# Dynamics of lipidome in a colon simulator

**DOI:** 10.1101/2022.12.13.520210

**Authors:** Matilda Kråkström, Alex M. Dickens, Marina Amaral Alves, Sofia D. Forssten, Arthur C. Ouwehand, Tuulia Hyötyläinen, Matej Orešič, Santosh Lamichhane

## Abstract

Current evidence suggests that gut microbiome derived lipids play crucial role in the regulation of host lipid metabolism. However, not much is known about the dynamics of gut microbial lipids within the distinct gut biogeographic. Here we employed targeted and untargeted lipidomics in the *in vitro* derived feces. Simulated intestinal chyme was collected from *in vitro* gut vessels (V1–V4), representing proximal to distal parts of the colon after 24 and 48 h with/without PDX treatment. In total 44 simulated chyme samples were collected from the *in vitro* colon simulator. Factor analysis showed that vessel and time had the strongest impact on the simulated intestinal chyme lipid profiles. We found that levels of phosphatidylcholines, sphingomyelins, triacylglycerols and endocannabinoids were altered in at least one vessel (V1–V4) during simulation. We also found that concentrations of triacylglycerols, diacylglycerols and endocannabinoids changed with time (24 vs. 48 h simulation). Together, we found that the simulated intestinal chyme revealed a wide range of lipids that remained altered in different compartments of the human colon model over time.

## 1. Introduction

Human gut harbors trillions of microbes that exhibit mutually beneficial relationship with the host [1]. A key contribution of the gut microbiota to the host is nutrient and xenobiotic metabolism, which plays a major role in training the immune system and promoting the intestinal homeostasis [2, 3]. Moreover, gut microbes are essential for the maintenance of the host metabolic homeostasis. Specific disturbances in the gut microbiome composition may contribute to wide-range of diseases, including inflammatory bowel disease [4], non-alcoholic liver disease [5] and psychiatric disorders [6]. Identifying the compositional changes in the gut microbiome alone, however, does not necessarily lead to mechanistic understanding [1]. During the last decade, metabolomics has emerged as a powerful approach in the microbiome field, providing functional information about the human gut microbial phenotype [7]. Therefore, a combined microbiome and metabolome strategy to evaluate host–microbiome interactions is being increasingly utilized.

To date, focus of the most metabolomics studies aimed at elucidation of the role of the gut– microbiome metabolome co-axis, have primarily been on water-soluble polar metabolites (e.g., tryptophan catabolites such as indole acetic acid, short-chain fatty acids (SCFA)). Other, non-polar microbial metabolites, including lipids such as sphingolipids (SLs), endocannabinoids (ECs), cholesterol, bile acids and acyl carnitines are less studied in comparison. However, lipids also have an important role in gut microbiome-host interactions [8]. Gut microbiota not only regulates intestinal lipid absorption and metabolism, but also impacts levels and metabolism of a substantial proportion of circulating lipids [9, 10]. Lipids are critical biomolecules involved in a wide range of cellular functions including, structure, communication and metabolism.

Lipidomics analysis of feces can identify numerous microbial lipids, which can inform about the gut microbial phenotype [11]. However, only a limited number of studies have integrated the lipidome in microbiome analyses with respect to health outcomes. In addition, dynamics of microbial lipids in the gut is poorly understood. This could be ascribed to the fact that it is not feasible to perform dynamic sampling across the human gastrointestinal tract. To overcome this challenge, *in vitro* colon models have been extensively applied to study the microbial functions [12]. Here we employed lipidomics approach on the *in vitro* derived intestinal chyme to examine the temporal lipid changes, occurring in different compartments of the colon simulator representing the proximal to the distal part of the colon. We also studied whether *in vitro* gut lipidome profiles were affected by colon simulation time and polydextrose (PDX), a synthetic complex oligosaccharide.

## 2. Materials and Methods

### 2.1 *In vitro* colon simulator

The Enteromix model of the human large intestine (Fig. 1) was described in detail previously [13, 14]. In summary, this simulator consists of eight separate units, each containing four semi-continuously fed connected glass vessels. The vessels in one unit (V1–V4) model the different compartments of the human colon from the proximal to the distal part, each having a different controlled pH and flow rate. Each unit is kept anaerobically and at 37 °C. In the initial phase of the simulation, each unit is inoculated with pre-incubated fecal microbes from a fresh fecal sample, which form the microbiota of the colonic model. In the present study, the fecal samples for inoculum were provided voluntarily by three healthy Finnish volunteers. The study and all methods used in it were carried out in accordance with relevant guidelines and regulations, and informed consent was orally obtained from all research subjects. This simulation was performed at IFF, Kantvik, Finland. To understand the lipidomic changes over time, the microbial slurry was collected from all vessels (V1–V4) after 24 and 48 h with/without PDX treatment. In total 44 samples were collected from the *in vitro* colon simulator vessels (V1–V4) and stored at −80 °C prior to lipidomic analysis. In addition, also media and inoculum used for the simulation and the pooled human fecal sample as a quality control sample were collected.

**Figure 1.**
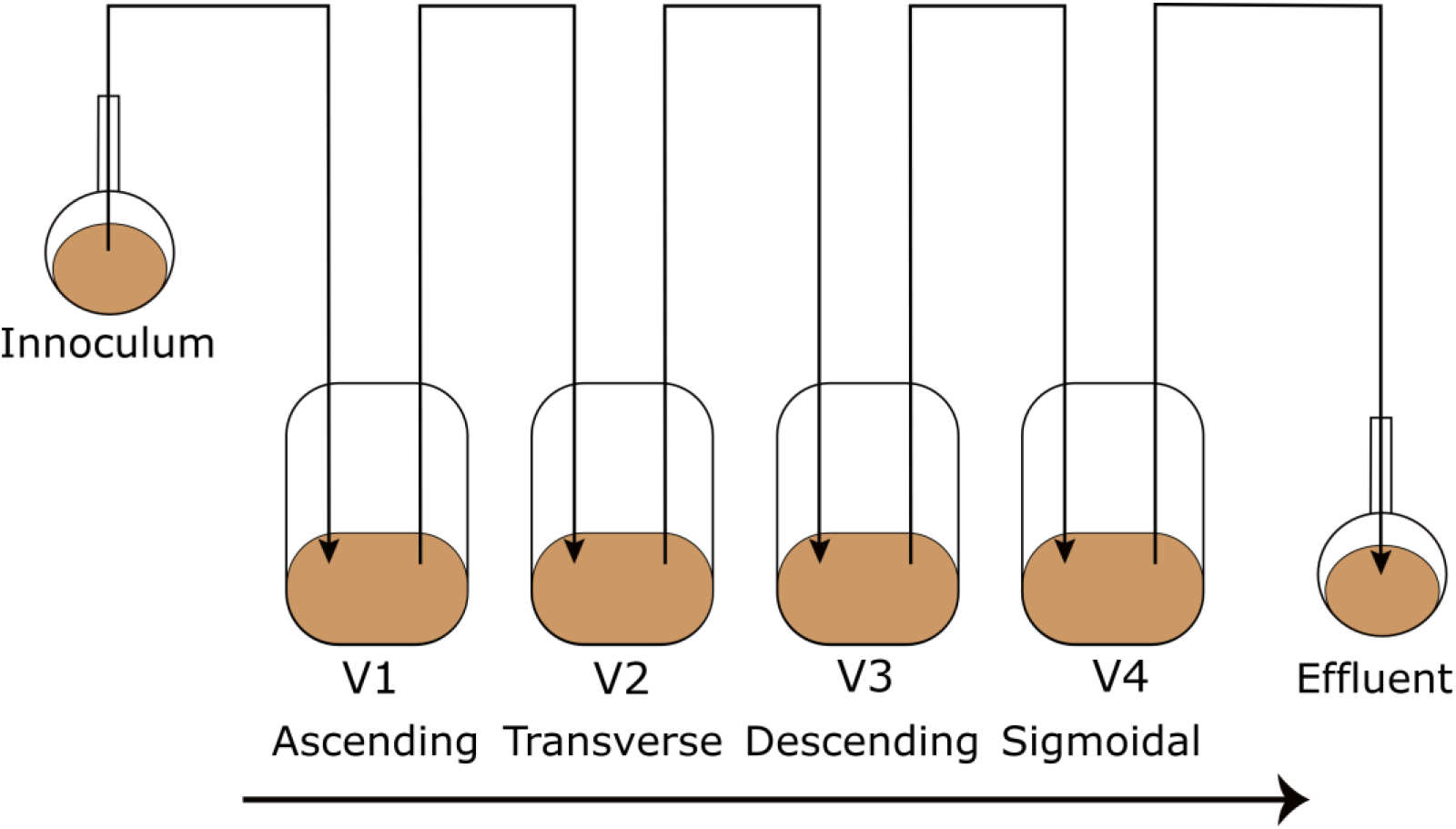
Schematic representation of the simulated colon model Enteromix. The vessels in one unit (V1–V4) simulate the different compartments of the human colon from proximal to the distal part, each having a different controlled pH and flow rate. The whole unit is kept anaerobically and at 37 °C. See [13] for more details.

### 2.2 Lipidomics analysis

Simulated intestinal chyme lipid extracts were prepared using a method based on the Folch procedure [15]as detailed by Lamichhane et al. [16]. An internal standard mixture containing 2.5 µg/mL mL 1,2-diheptadecanoyl-sn-glycero-3-phosphoethanolamine (PE(17:0/17:0)), N-heptadecanoyl-D-erythro-sphingosylphosphorylcholine (SM(d18:1/17:0)), N-heptadecanoyl-D-erythro-sphingosine (Cer(d18:1/17:0)), 1,2-diheptadecanoyl-sn-glycero-3-phosphocholine (PC(17:0/17:0)), 1-heptadecanoyl-2-hydroxy-sn-glycero-3-phosphocholine (LPC(17:0)), 1-palmitoyl-d31-2-oleoyl-sn-glycero-3-phosphocholine (PC(16:0/d31/18:1)) and 1,2,3-triheptadecanoyl-sn-glycerol (TG(17:0/17:0/17:0)) was prepared in CHCl_3_:MeOH (2:1, v/v). Six point calibration curves with concentrations between 100 and 2500 ppb in CHCl_3_:MeOH (2:1, v/v) were prepared for 1-Hexadecanoyl-2-octadecanoyl-sn-glycero-3-oethanolamine (PE(16:0/18:1)), octadecenoyl-sn-glycero-3-phosphocholine (LPC(18:1)), Cholesteryl hexadecanoate (CE(16:0)), 1,2-Distearoyl-sn-glycero-3-phosphoethanolamine (PE(18:0/18:0)), N-stearoyl-D-erythro-sphingosylphosphorylcholine (SM(18:0/18:1)) and Cholesteryl linoleic acid (CE(18:2)). The samples were prepared by spiking 10 µL of sample with 10 µL of 0.9% NaCl and 120 µL of internal standard solution. The samples were vortexed and were left to stand on ice for 30 min. Samples were centrifuged (9400 × g, 5 min, 4 °C) and 60 µL from the lower layer was diluted with 60 µL of CHCl_3_:MeOH (2:1, v/v). For the LC separation, a Bruker Elute UHPLC system (Druker Daltonik, Bremen, Germany) equipped with an auto sampler cooled to 10 °C, a column compartment heated to 50 °C and a binary pump was used. A Waters ACQUITY BEH C18 column (2.1 mm × 100 mm, 1.7 µm) was used for chromatographic separation. The flow rate was 0.4 mL/min and the injection volume was 1 µL. The needle was washed with 10% DCM in MeOH and ACN: MeOH: IPA: H_2_O (1:1:1:1, v/v/v/v) + 0.1% HCOOH after each injection for 7.5 s each. The eluents were H_2_O + 1% NH_4_Ac (1M) + 0.1% HCOOH (A) and ACN: IPA (1:1, v/v) + 1% NH_4_Ac + 0.1% HCOOH (B) The gradient is as follows: from 0 to 2 min 35-80% B, from 2 to 7 min 80-100% B and from 7 to 14 min 100% B. Each run was followed by a 7 min re-equilibration period under initial conditions (35% B).

Mass spectrometric detection was performed on a Bruker Impact II QTOF (manufacturer, city, country). For data pre-processing, the raw data files were converted to .mzml file using Bruker compass data analysis (version no). The pre-processing was performed in mzmine2 according to (reference). Briefly, centroid mass detection was performed, followed by ADAP chromatogram builder, chromatogram deconvolution (local minimum search), isotopic peak grouper a join aligner. After this, a filtering step (Feature list rows filter), a custom database search, an adduct search and gap-filling (Peak finder) was performed. Finally, the results were exported as a csv file. After this, lipid class-based normalization was performed using the class-based internal standards, class-based calibration curves were created, and semi-quantification was performed using the calibration curves. Features which were annotated and had a relative standard deviation of less than 30% in the QC samples were selected for further processing.

### 2.3 Endocannabinoid analysis

Crash solvent (400 µL) consisting of acetonitrile (ACN), 0.1% formic acid (FA) and isotopically labelled internal standards (Supplementary Table S1) was added to a glass vial and 200 µL of chyme slurry was added. The samples were vortexed and left to settle at -20 °C for 30 minutes. The samples were filtered through protein precipitation filter plates and collected into 96 well plates with glass inserts. The samples were transferred to glass vials and dried at 35 °C under a gentle stream of nitrogen. The samples were reconstituted in 50 µL reconstitution solution (60% water, 20% ACN and 20% isopropanol). The samples were analyzed using LC-MS. The chromatographic separation was performed on a Sciex exion (AB Sciex Inc., Framingham, **MA)** consisting of a binary pump, an autosampler and a thermostated column compartment. The column used was an XBridge BEH C18 2.5µm, 2.1×150mm column with a pre-column made with the same material. The eluents were A: 0.1 % FA and 1% ammonium acetate (1M) in water and B: 0.1 % FA and 1 % ammonium acetate (1M) in ACN/IPA (50:50). The gradient is presented in Supplementary Table 2, the injection volume was 1 µL, the flow rate was 0.4 mL/min and the column oven temperature was 40 °C. The detection was performed on a Sciex 7500 QTrap operating in MRM mode. The parameters used are presented in Supplementary Table 1 and 3. Quantification was performed using calibration curves from 0.01 ppb to 80 ppb (0.1 and 800 ppb for Arachidonic acid (AA)) using the internal standard method. Quantification was performed with Sciex OS analytics.

### 2.4 Data analysis

Lipid data values were log-transformed prior to multivariate analysis. The difference in the lipidome between different vessel, time and case (with/without PDX treatment) were analyzed using a multivariate linear model using MaAsLin2 package in R (lipids ∼ Time + Vessel+ case). Adjusted p-values with FDR = 0.25 were considered significant. Spearman correlation coefficients were calculated using the Statistical Toolbox in MATLAB 2017b and p-values < 0.05 (two-tailed) were considered significant for the correlations. The individual Spearman correlation coefficients (R) were illustrated as a heat map using the ‘‘corrplot’’ package (version 0.84) for the R statistical programming language.

## 3. Results

### 3.1. Untargeted lipidomics and targeted endocannabinoid analysis in the simulated faecal samples

We analyzed simulated intestinal chyme lipids obtained from different vessels (V1-V4) in the in vitro colon, which mimicks the compartments of the human colon from the proximal to the distal part (Figure 1). In total, 44 simulated intestinal chyme samples were collected from the in vitro colon simulator (vessels V1–V4) ran for either 24 or 48 hours with/without PDX treatment. The untargeted lipidomics assay of the simulated chyme extract resulted in the detection of 118 annotated lipids. These lipids were semi-quantified using class-specific internal standards and calibration curves. The semi-quantified lipids included a wide range of lipid classes: triacylglycerols (TG), ceramides (Cer), cholesterol esters (CE), diacylglycerols (DG), lysophosphatidylcholines (LPC), phosphatidylcholines (PC), phosphatidylethanolamines (PE), and sphingomyelins (SM). Of the 13 endocannabinoids (EC) studied, 11 were detected in at least one sample type. These included Palmitoyl ethanolamide (PEA), Arachidonoyl glycerol (AG), 2-Arachidonic glycerol ether (2-Age), Arachidonoyl ethanolamide (AEA), Oleoyl ethanolamide (OEA), Stearoyl ethanolamide (SEA), Docosatetraenoyl ethanolamide (DEA), Alpha-linolenoyl ethanolamide (aLEA), Arachidonic acid (AA), N-arachidonoyl taurine (NAT) and N-arachidonoyl-L-serine (NALS). Nine (PEA, AG, 2-AGe, AEA, OEA, SEA, DEA, aLEA and AA) were detected in human faeces, which were used as quality control samples.

### 3.2 Lipidome in the in vitro colon simulator

Principal Components Analysis (PCA) of the preprocessed lipidomics data revealed a clear vessel-related pattern in the simulated intestinal chyme samples. To examine the contributions of various factors to simulated chyme lipidome profiles, multivariate linear modelling was perfomed (lipids ∼ Time + Vessel + treatment). We found that vessel had a marked impact on the simulated chyme lipidome, when compared to the simulation time (24 vs 48 h) and treatment (with/without PDX treatment). Of the analyzed lipids, 40 showed a significant change in at least one of the vessel (p < 0.05, Figure 2A and Supplementary Table S4). These lipids included one CE, two DGs, five Cer, eleven PCs, one PG, four PEs, four SMs and nine TGs (Figure 2A-C). All of these lipids passed the FDR threshold of 0.25 (Supplementary Table 1). With the exception of Cer(d18:1/20:0), LPE (16:0), LysoPE(18:1), most of the lipids showed a decreased pattern from vessel V1 to V4, i.e. from the proximal to the distal part of the colon simulator (Figure 2B-D). Among the TGs, specifically, those TGs with low double bond count (≤ 2 double bonds) showed changes within the vessels (V1-V4, Supplementary Table S4). However, no clear pattern with respect to the double-bond counts and/or carbon numbers composition was observed in any other specific lipid class.

**Figure 2.**
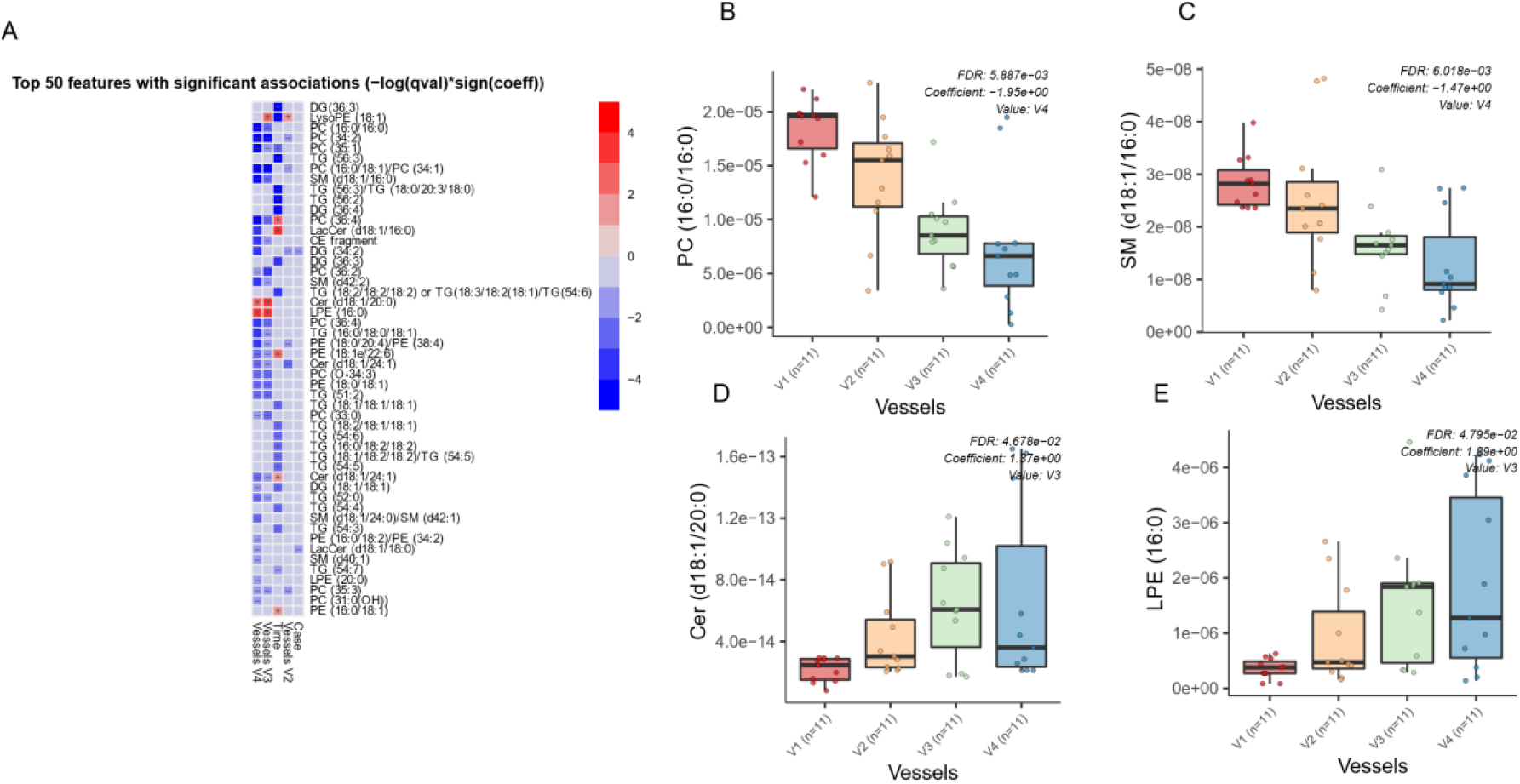
Dynamics of lipids in the in vitro colon simulator. The figure depicts the lipidomic changes in four vessels (V1–V4) over two time points (24 and 48 h). A) Heat map showing the difference in the lipidome between vessels (V1-V4), time (24 vs 48) and case (with/without PDX) compared using multivariable linear model. The changes in vessels were reference to vessel 1. Blue indicate decreasing trend while red denotes increase in the trend. C-D Box plot shows the difference in the lipidome between vessels (V1-V4). These selected lipids highlights the trend between the vessels (V1 to V4), which mimicks the compartments of the human colon from the proximal to the distal part We also found that chyme lipidome concentrations of 26 lipids, mainly TGs (n = 14) were different across two time points (24 vs. 48 h of simulation, Supplementary Table S5), while only four lipids (DG(34:2), LacCer(d18:1/18:0), TG (55:6) / TG (16:0/19:1/20:5), TG (51:1)) were found altered when simulation was performed with/without PDX.

Next, we analyzed the dynamics of ECs in different compartments of the colon simulator over time. Among the 11 detected ECs, the level of 7 ECs were altered in at least one of the vessel in the colon simulator (p < 0.05, Figure 3A and Supplementary Table S6). These ECs include AEA, aLEA, DEA, OEA, PEA, and SEA. There was no persistent trend, but the level of OEA, aLEA and PEA increased in vessel (V3–V4) when compared to vesssel V1 (Figure 3B). A similar trend was seen for AEA with higher level of variation appearing in the vessel V3 (Figure 3C). On the other hand, the level of SEA was increased in V1 when compared to vessels V2-V4 (Figure 3 and Supplementary Figure S1). In addition, we analyzed the EC concentrations over time in the simuated gut. Overall, five ECs (AEA, 2-AGe, AA, PEA, and NAT) were increased in the colon simulation ran for 48 h when compared to 24 h intestinal chyme slurry (Supplementary Figure S2).

**Figure 3.**
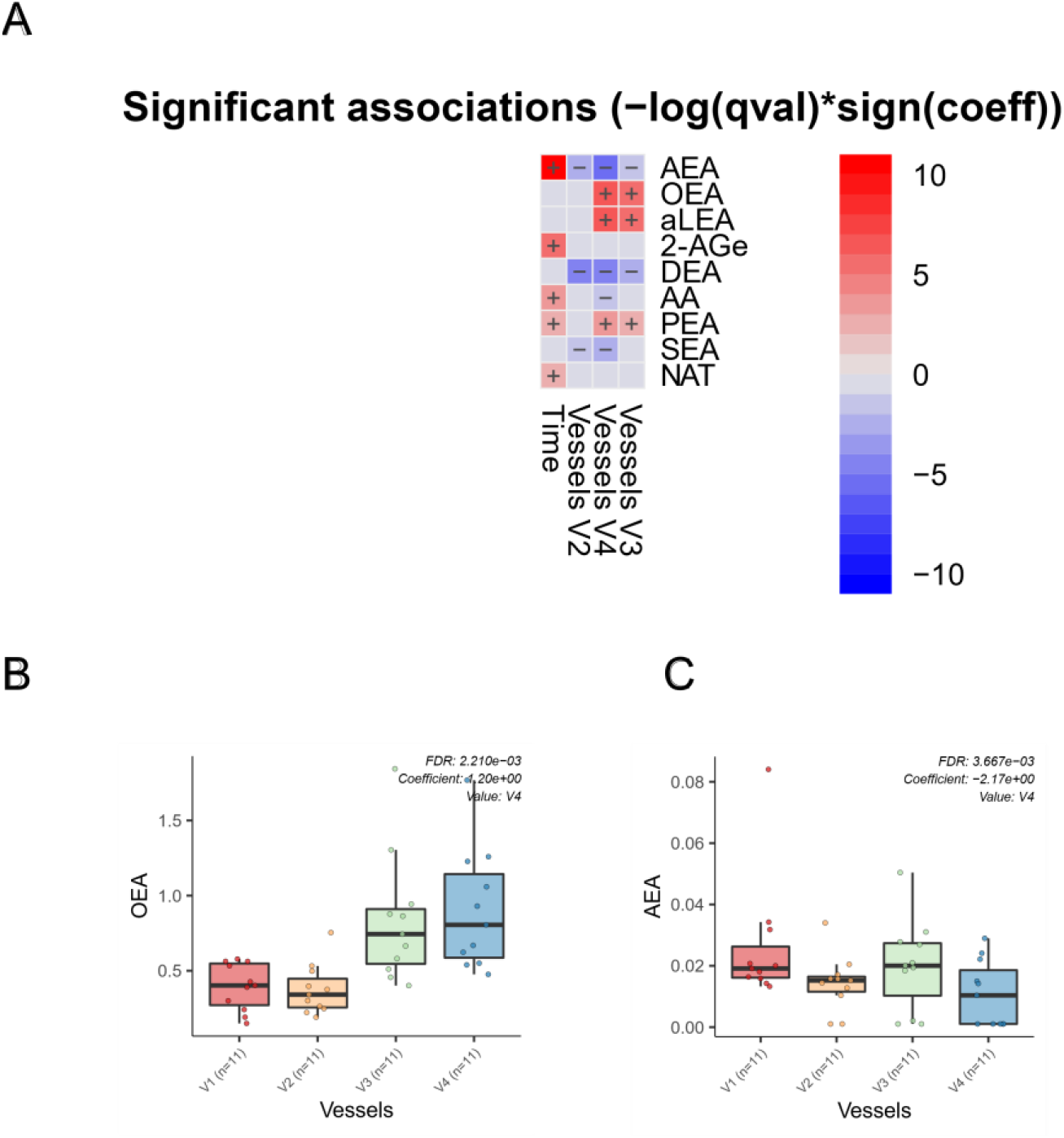
Dynamics of ECs in the in vitro colon simulator. The figure depicts the EC changes in four vessels (V1–V4) over two time points (24 and 48 h).A) Heat map showing the difference in the EC between vessels (V1-V4), time (24 vs 48) and case (with/without PDX) compared using multivariable linear model. The changes in vessels were reference to vessel 1. Blue indicate decreasing trend while red denotes increase in the trend. B-C Box plot shows the difference in the EC between vessels (V1-V4). These selected EC highlights the dynamics trend between the vessels (V1 to V4), which mimicks the compartments of the human colon from the proximal to the distal part.

Given the known link between gut microbiota and EC metabolism [17], we also examined difference in ECs profiles between the media used for in vitro gut simulation and the simulated chyme slurry. We found most of the ECs dectected in the simulated chyme slurry were lower in concentration than in the simulation media. Interestingly, we observed NALS was detected at low concentrations in the media, however, it was not detected in any of the vessels (Figure 4A). Meanwhile, 2-AGe was not detected in the media, but appeared in different vessels (Figure 4B, V1-V3).

**Figure 4.**
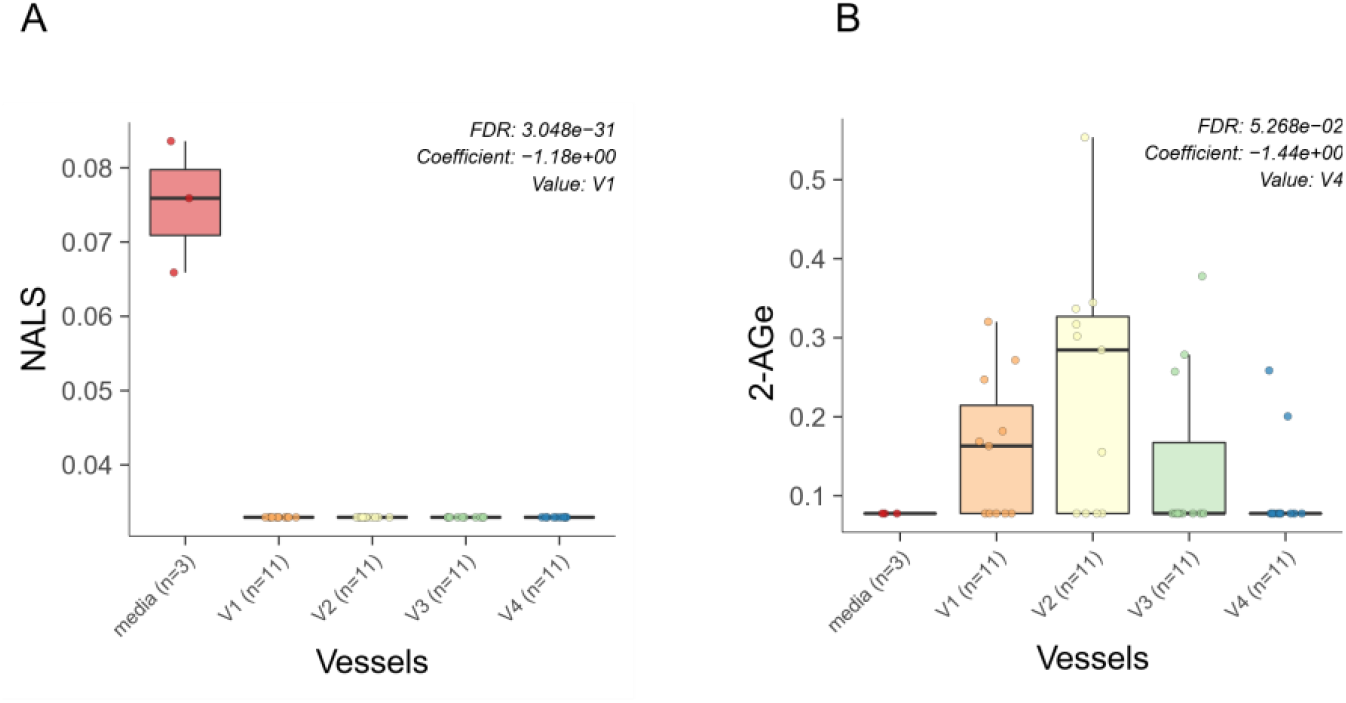
ECs in the media vs. in vitro colon simulator. The figure depicts the endocannabinoid changes in four vessels (V1–V4) over two time points (24 and 48 h).A-B Box plot shows the difference in the EC between vessels (V1-V4) and media used during the simulation.

### 3.3. Association of lipidome and ECs in the in vitro colon simulator

Next, we performed correlation analysis between ECs and individual simulated intestinal chyme lipid levels (Figure 5). We found the levels of SEA and DEA were positively associated with the overall simulated chyme lipidome. OEA, aLEA and PEA showed positive association with Cers and LPEs, while those being inversly correlated with PCs, PEs, SMs and TGs. This trend was less pronounced for AA and 2-AG. Instead, there was a clear, inverse association trend between SMs, and PEs and AA level in the in vitro simulator. In addtion, DGs were negatively related with the level of AA and 2-AG. No association between individual lipids and NAT was observed, with exception for a Cer(d18:1/20:0).

**Figure 5.**
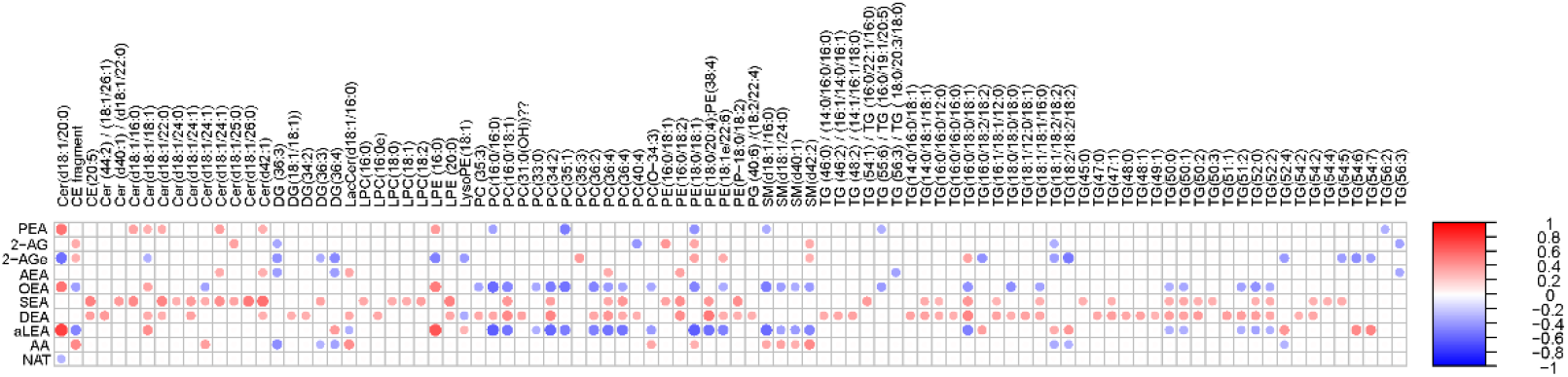
Association of lipidome and ECs in the in vitro colon simulator. Spearman correlation coefficients illustrated by heat map.

## 4. Discussion

In this study, we reported the dynamics of lipids in each compartment of the colon simulator. We found a more distinct lipids profile in the proximal colon (vessel V1) than in the distal part of the colon (vessel V4). Specifically, specific Cer, PCs, SMs and TGs were decreased in the distal as compared to the proximal part of the colon simulator. Our observations also show that *in vitro* derived intestinal chyme lipids, particularly TGs, are strongly affected by time. Our results are in agreement with previous studies, showing that profound metabolic changes occur in different parts of the *in vitro* gut over time [12]. The level of SCFAs (acetate, butyrate, and propionate), branched chain fatty acids (iso-valerate), biogenic amines (trimethylamine), organic metabolites (succinate, ethanol, formate, valerate, and n-acetyl compound) and amino acids (lysine, leucine, iso-leucine, phenylalanine, tyrosine, and valine) were reported to change in the four vessels (V1–V4) within 48 h (12, 24, 36, and 48 h) [12, 14]. Similar dynamic changes in metabolome along the intestinal tract has been reported in the nonhuman primates [18].

The human gut is an endogenous source of systemic sphingolipids [19]. We observed distinct changes in the levels of the sphingolipids (Cer and SMs) while passing from vessel (V1–V4) over time (24 and 48 h). Sphingolipids were higher in the proximal colon (V1) than in the distal part of the colon (V4). Sphingolipids have many structural and signaling roles in eukaryotes [19] and microbially-derived sphingolipids may markedly impact the host sphingolipid levels [20, 21]. In addition, sphingolipids have been shown to promote the survival of commensal bacteria [22]. Notably, the observed decreasing trend in our study may indicate increased microbial catabolism. Lipid is considered an alternative pathway for carbon, nitrogen, and an energy source for the gut microbes [23]. Microbiome data is not available in our study, therefore, we could not demonstrate causal link between gut microbiota and sphingolipid metabolism. Notwithstanding this, our study introduced a simulated intestinal chyme lipidomics-based method to interrogate potential microbial lipids within the complex gut system. This study also highlights that an *in vitro* colon model provides a controlled experimental condition, i.e., dynamic sampling for characterization of the gut microbiota-derived bioactive lipids.

Growing evidence suggest that bidirectional interactions exist between gut microbes and the endocannabinoid system [24-27]. The endocannabinoid system comprises network of cannabinoid type receptor and its ligand (i.e., the endocannabinoids) that are present throughout the human body. The concept of endocannabinoid system and the gastrointestinal tract is over a half century old. Studies from the 1970s already showed that ECs can have profound impact on gut motility [17, 28, 29]. Current evidence suggests that gut transit time affects the gut microbial composition and function [30]. Here we found that concentration of ECs varied between proximal (V1) and distal colon (V4). We also found an endocannabinoid of potential microbial origin (i.e., 2-AGe). Phospholipids are precursors of ECs [31]. Intriguingly, we found inverse association between PCs and those ECs that were increasing from V1 to V4. We thus hypothesize that availability of PCs as substrate in the gut drives the level of specific ECs. In addition, this finding highlights the role of microbes in metabolism of ECs in the gut.

In summary, our study shows that the combination of lipidomics and *in vitro* derived intestinal chyme samples enabled us to characterize the fate of lipids in a simulated human colon. Our study also reports novel profile of ECs in the simulated intestinal chyme.

## Supplementary Materials

The following supporting information can be downloaded at: www.mdpi.com/xxx/s1, Figure S1: SEA in four vessels (V1–V4); Figure *S*2: ECCs in four vessels (V1–V4) over time (24 hours vs 48 hours). Table S1-S6.

## Author Contributions

“Conceptualization, S.L. M.O., A.D., and T.H.; methodology, M.K., A.D., and M.A.A.; software, formal analysis, M.K. and S.L.; investigation, M.K. and S.L.; resources, S.D. F. and A.C.O; data curation, M.K., A.D., and M.A.A.; writing—original draft preparation, S.L. and M.K; writing—review and editing, all authors.; visualization, S.L.; supervision, S.L. M.O., A.D., and T.H.; acquisition, M.K., and M.A.A.;. All authors have read and agreed to the published version of the manuscript.”

## Funding

This work was supported by Academy of Finland (no. 323171 to S.L.), Academy of Finland (No. 333981 to M.O.), Novo Nordisk Foundation (NNF19OC0057418 to M.O.), and Swedish Research Council (no. 2016-05176 to T.H., no. 2018-02629 to M.O.).

## Informed Consent Statement

The fecal samples used as inoculum in the colon simulator were given voluntarily by healthy adult Finnish volunteers. According to the Finnish law, no ethical approval is needed for this kind of study, since there has not been any interference with a person’s privacy, physical or mental integrity. “The study presented in the report is not considered medical research as defined in the Finnish Act on Medical Research (488/1999, as amended). Due to this, the study did not require an approval from the ethical committee and therefore such approval has not been obtained. In addition, as the study is not considered as medical research, the consent has not been obtained in writing as required by the Act on Medical research, but orally.”. This is a legal statement, and should not be rephrased/rewritten.

## Data Availability Statement

The lipidomics datasets generated in this study will be submitted to the Metabolomics Workbench repository (https://www.metabolomicsworkbench.org).

## Acknowledgments

We thank the Turku Metabolomics Center for the assistance and resources in the analyses of fecal ECCs and lipidome.

## Conflicts of Interest

Sofia D. Forssten and Arthur C. Ouwehand are employees of Danisco Sweeteners Oy, IFF Health & Biosciences (Kantvik, Finland).

